# Social networks with strong spatial embedding generate non-standard epidemic dynamics driven by higher-order clustering

**DOI:** 10.1101/714006

**Authors:** David J. Haw, Rachael Pung, Jonathan M. Read, Steven Riley

## Abstract

Some directly transmitted human pathogens such as influenza and measles generate sustained exponential growth in incidence, and have a high peak incidence consistent with the rapid depletion of susceptible individuals. Many do not. While a prolonged exponential phase typically arises in traditional disease-dynamic models, current quantitative descriptions of non-standard epidemic profiles are either abstract, phenomenological or rely on highly skewed offspring distributions in network models. Here, we create large socio-spatial networks to represent contact behaviour using human population density data, a previously developed fitting algorithm, and gravity-like mobility kernels. We define a basic reproductive number *R*_0_ for this system analogous to that used for compartmental models. Controlling for *R*_0_, we then explore networks with a household-workplace structure in which between-household contacts can be formed with varying degrees of spatial correlation, determined by a single parameter from the gravity-like kernel. By varying this single parameter and simulating epidemic spread, we are able to identify how more frequent local movement can lead to strong spatial correlation and thus induce sub-exponential outbreak dynamics with lower, later epidemic peaks. Also, the ratio of peak height to final size was much smaller when movement was highly spatially correlated. We investigate the topological properties of our networks via a generalized clustering coefficient that extends beyond immediate neighbourhoods, identifying very strong correlations between 4th order clustering and non-standard epidemic dynamics. Our results motivate the joint observation of incidence and socio-spatial human behaviour during epidemics that exhibit non-standard incidence patterns.

**Author Summary:** Epidemics are typically described using a standard set of mathematical models that do not capture social interactions or the way those interactions are determined by geography. Here we propose a model that can reflect social networks influenced strongly by the way people travel and we show that they lead to very different epidemic profiles. This type of model will likely be useful for forecasting.

## Introduction

Epidemics are frequently conceptualized as resulting from the transmission of a pathogen across a network. Directly transmitted pathogens propagate through susceptible human populations and create directed infection trees with an offspring-like process [15]. Each node may be a different type (e.g. children may be more infectious than adults [42]) and individuals with many contacts are more likely to cause infection than those with fewer contacts [24]. Although difficult to observe, infection trees describe a real biological process: these pathogens do not reproduce outside of a human host, so the founding pathogen population for an infectee comes directly from their infector. Further, we can conceptualise that infection trees occur when a true offspring process is constrained to pass through a social network [3, 40], with infection occurring according to a specified probability when an edge exists between a susceptible and an infectious individual.

The properties of different contact network types can be described by distributions associated with their topology [40]. First order network properties are associated with first order connections, as defined by the degree distribution. For finite random networks of reasonable size, the degree distribution is well-approximated by a Poisson in which variance is equal to the square of the mean. In contrast, for finite scale-free networks, the offspring distribution is power law-like with a much higher variance. Further, distributions of second order phenomena describe connections of length two. For example, the local clustering coefficient is a second order property, defined to be the neighbourhood density of a given node [40]. For a limited set of network types, we can use analytical expressions for higher moments of the degree distribution to calculate key properties of their potential epidemics, such as the probability of epidemic establishment and cumulative incidence [22, 28]. Although these higher order moments are tractable for some special cases, they are seldom the primary target of theoretical studies. Semi-empirical networks that arise from detailed simulations [11] may have complex higher moments, however their impact on epidemic dynamics is obscured by the variance of their offspring distribution e.g. [25]. Here, we explicitly control our network generation algorithm so as to have non-trivial higher order structure whilst maintaining a Poisson degree distribution and a pre-specified clustering coefficient.

Epidemics can also be understood in terms of compartmental models, which are more tractable mathematically, and are equivalent to large network models with very simple topologies [35]. Key features of epidemic incidence curves is are often explained by dynamics associated with these models [1, 19]. Numerical solutions to multi-type SIR-like compartmental models are easier to obtain than for many topologies of network and can explain: the initial growth phase [30], the timing and amplitude of the peak [43], epidemic duration [21] and the total number of cases [17]. These models can efficiently describe many different types of complexity, such as age-specific susceptibility and transmissibility [16], behavioural risk groups [4] and, with increasing frequency, geographical location [34].

The basic reproductive number has been defined for both compartmental models and for network models. For compartmental models, the reproduction number is conditional on the system having a well defined period of exponential growth [18] and is defined as the average number of new infections generated by a typically infectious individual in an otherwise infectious population [18]. The word “typically” is somewhat overloaded in this definition: during the exponential phase, a system with heterogeneous population will reach a steady-state distribution of infectives, corresponding to the eigenstate of the renewal process.

For network models, the basic reproduction number is most frequently defined as the expected ratio of cases between the first (seed) and second generations of infection. In homogeneous networks, this is equal to the product of the average degree and the probability of transmission per link per generation. However, many studies of epidemics on networks involve high variance degree distributions [26, 25], and so this quantity must be modified to account for excess degree [26, 27]. Here, we use *R*^∗^ to denote the expected first generation ratio *if a network is homogeneous*, defined to be the expected number of cases in the second generation divided by the number in the first generation. Our *R*^∗^ is therefore consistent with *ρ*_0_ as defined in [26], although we choose not to adjust for over-dispersion, because we condition our network construction on this distribution having low variance.

The reproduction number for networks has also been defined to be more consistent with its definition for compartmental models. In [37] *R*_∗_ was defined as an asymptotic property of epidemics that were guaranteed to have an exponential phase when they occurred on infinitely large networks. We define our *R*_0_ to be a finite-network approximation to this *R*_∗_ in [37]. This *R*_0_ is well-defined during periods of exponential growth.

Both compartmental and network models can be embedded in space [34]. Each node can have a location in space while each compartment can refer to a single unit of space. Node density can be assigned according to known population densities and compartments can be assigned equal spatial areas but different numbers of hosts. In general, the risk of infection passing between two people decreases as the distance between their home location increases. The propensity of nodes to form links across space or for infection to spread between compartments can be quantified using mobility models borrowed from geography [12], such as the gravity and radiation models. Here, we are specifically interested in how the overall topology of a spatially-embedded network model can be driven by different movement assumptions and thus drive the gross features of the epidemics that occur on the network.

## Results

We used an existing variant of the Metropolis-Hastings algorithm [35] to create a spatially-embedded bipartite network of homes and workplaces consistent with the population density of Monrovia, Liberia, and with three illustrative movement scenarios (SI Appendix, Fig S1). An individual’s propensity to choose a given workplace was determined by the distance between their home and workplace and parameters of a gravity-like kernel. The kernel was inversely proportional to distance raised to the power *α*, with movement scenarios generated solely by changing the value of *α*: a control value *α* = 0 that removed the embedding and produced a *non-spatial* model; a *wide* kernel with *α* = 3 typical of developed populations [35, 38]; and a *highly local* kernel with *α* = 6 representing less developed populations (SI Appendix, Fig S1 part C compared with rural Huangshan in Ref [14]). The resulting distributions of distances from home to work were driven strongly by our choice of *α*, with 95% of journeys: less than 24.12km for *α* = 0; less than 12.91km for *α* = 3; and less than 6.68km for *α* = 6. Workplace links were dissolved into links between individuals in different households resulting in a network of cliques (households) that were linked according to *α*.

The choice of movement kernel used to create the household-workplace networks affected gross features of simulated epidemics, even when controlling for other aspects of the network topology (Fig 1). Unipartite contact networks between households were obtained from the bipartite network of households and workplaces and were dependent on three parameters: mean household size *h*, mean number of workplace links *v*, and probability of forming a link in the work-place *p_w_*. The mean workplace size *w* and mean degree of the network were determined by these parameters: *w* = *v/p*_*w*_ + 1, ⟨*k*⟩ = *h* − 1 + *v*. Across a broad range of plausible values for *h*, *v* and *p_w_*, very local movement (*α* = 6) produced later epidemics than did typical developed-population movement (*α* = 3) or spatially random mixing (*α* = 0, Fig 1A). Similarly, time to extinction was later for very local movement (*α* = 6) compared with more frequent longer-distance movement (*α* = 3) or the absence of spatial embedding (*α* = 0). We calculate the coefficient of variation of the degree distribution 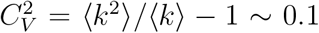 for each network, independently of *α* [26].

**Figure 1:**
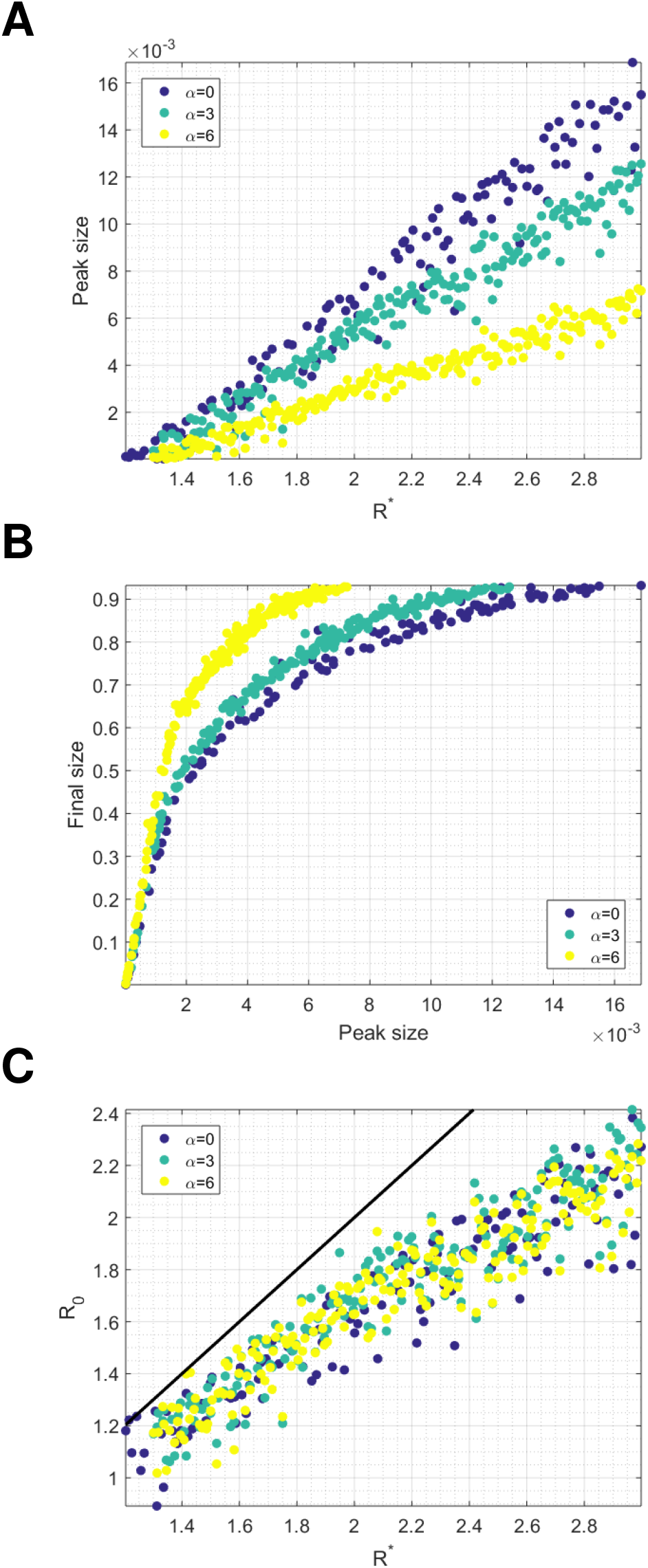
For each set of parameters drawn from the Latin hypercube, and for *α* = 0, 3, 6, we show relationships between: **(A)** *R*^∗^ and peak size, **(B)** peak size and final size, **(C)** *R*^∗^ and *R*_0_ (with the line *R*_0_ = *R*^∗^ shown in black).

Each simulation is assigned a value of *R*^∗^, the average number of cases in the first generation per seed infection. For moderate-to-high values of the first generation ratio *R*^∗^, there was very little difference in the final size of the outbreak for the different movement assumptions. However, for low values of *R*^∗^ < 1.8, the average final size of the outbreak was substantially smaller for more local kernels. This was driven by a higher probability of extinction when more local movement was assumed. The difference in final size driven by *α* was no longer present when we controlled for extinction (SI Appendix, Fig S2).

The choice of movement scenario had a substantial impact on peak incidence, even when *R*^∗^ was high and there was little difference in the final sizes (Fig 1B, Fig 2 rows 1 and 2). For example, for parameters with first generation ratios in the range [1.8, 2.2], average peak daily incidence as a fraction of the total population was 6.5 × 10^−3^ for random spatial movement, 5.4 × 10^−3^ for movement assumptions typical of developed populations and 3.0 × 10^−3^ when highly local movement was assumed. The relationship between peak height and first generation ratio appeared to be strongly linear, with correlation coefficients 0.9778, 0.9826 and 0.9806 for *α* = 0, 3 and 6 respectively.

**Figure 2:**
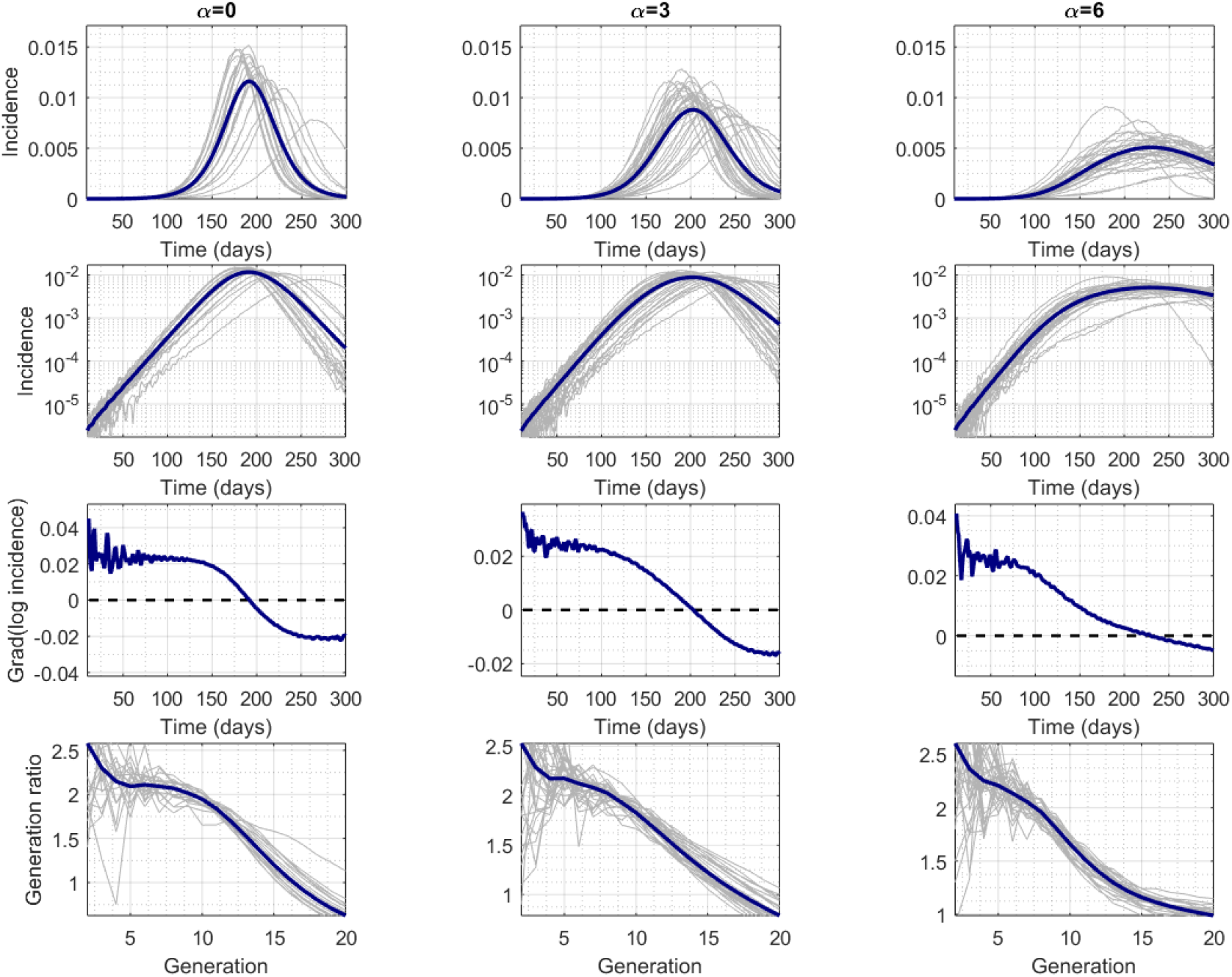
Columns correspond to network structures with *α* = 0, 3 and 6 and simulations with *R*_0_ ∈ (2, 2.2]. Exponential growth in real time is indicated by straight lines (second row) and horizontal lines (third row); horizontal lines in bottom row indicate exponential growth by generation. Figures S3 to S5 show results for a wider range of *R*_0_ values for *α* = 0, 3, 6.

The relationship between peak incidence and final size for the three movement scenarios illustrates further how clustering within the network directly affects gross features of an epidemic. Peak incidence is observed prior to final size during an epidemic. For the same peak height, local movement gave substantially larger final sizes. For peak daily incidences in the range [3 × 10^−3^, 6.5 × 10^−3^] the final size of the outbreak was 68% when random spatial movement was assumed, 74% when movement was assumed to be typical of developed populations and 84% when highly-local movement was assumed.

For all movement scenarios, the basic reproductive number *R*_0_ was smaller than the first generation ratio *R*^∗^ and different from the expected number of secondary cases generated by a single seed in an otherwise susceptible population. The duration of the exponential phase can be seen when incidence is plotted on a log scale: a constant gradient of log incidence is evidence of exponential growth (Figure 2, third row). However, in a network model with clearly defined generations, the generation ratio can also be used to define exponential growth: if the ratio of incidence between generation *n* + 1 and *n* is the same as the ratio between generations *n* and *n* − 1, then we can claim to have identified a period of exponential growth (Methods, Fig 2). The value of that constant observed ratio is the basic reproductive number *R*_0_ [18].

Incidence grew exponentially for a much shorter time for highly-local movement than it did for a wider movement kernel, or for non-spatial networks, even when we controlled for *R*_0_ to be within a narrow range (e.g. (2, 2.2], Fig 2). Despite this being a relatively large population, there was no obvious period of exponential growth when we assumed highly local movement. Therefore, given that the basic reproductive number is defined for a genuine renewal process – and its implied exponential growth [18] – it could be argued that *R*_0_ does not exist for some of these networks for our model parameters. However, we did assign a value of *R*_0_ for all simulations based on the most similar subset of consecutive early generations (see Methods). The amplitude of the difference was not driven in any obvious way by the underlying assumptions used to create the networks. These patterns were not specific to the range of values for *R*_0_ (SI Appendix, Figs. S3, S4, S5).

Analysis of the higher-order structure of the networks suggests that movement scenarios were driving the observed characteristics of epidemics such as peak timing and attack rate via increased fourth order clustering. We use the term first order clustering for the quantity typically described as the local clustering coefficient [40]: the link density of the immediate neighbour-hood of a given node. By extension, we defined order-*m* clustering coefficient to be the expected proportion of neighbours within *m* steps on the network who were also neighbours of each other within *m* steps (Fig 3). We found no relationship between our assumed pattern of movement (*α*) and first or second order clustering coefficients. There was a weak relationship between *α* and third order clustering and then a very strong relationship between *α* and forth order clustering. Patterns between epidemic properties and fourth order clustering for individuals were similar to those between epidemic properties and second order clustering of households, as would be expected, given the bipartite algorithm used to create individual-level networks.

**Figure 3:**
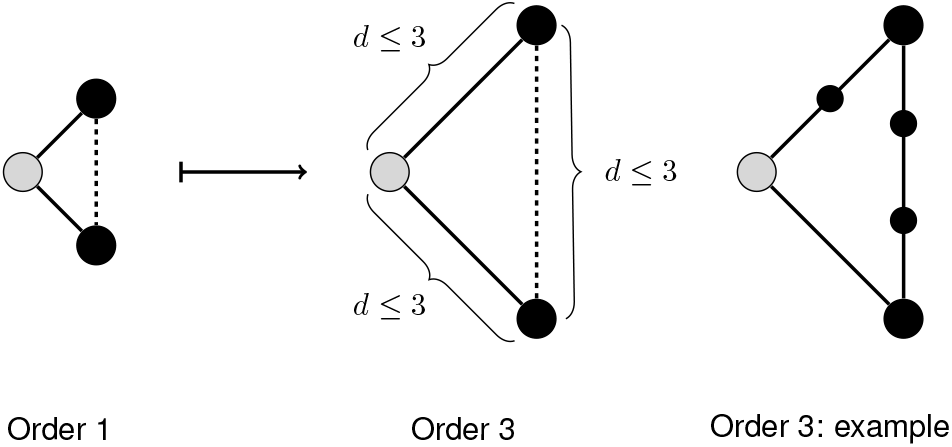
A schematic showing the generalization of clustering coefficient *CC*^1^ to higher orders 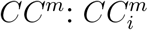 measures the density of paths of length *d* ≤ *m* between the up-to-*m* neighbours of node *i* (where node *i* is shown in gray).

Final size increased with spatial correlation, despite peak size displaying the opposite trend for controlled *R*^∗^ or *R*_0_. There was a strong linear relationship between order-*m* clustering and peak size/final size, that could be explained by *α*, the strength of spatial embedding, when we control for *R*_0_ (Fig 4B). The gradient of the relationship decreased with order of clustering. Second order household clustering showed the same relationship with peak size as did fourth order individual clustering (Fig 4C). These strong linear relationships only existed when we effectively control for *R*_0_, rather than *R*^∗^, and became less noisy when we reduced the interval used to define *R*_0_.

**Figure 4:**
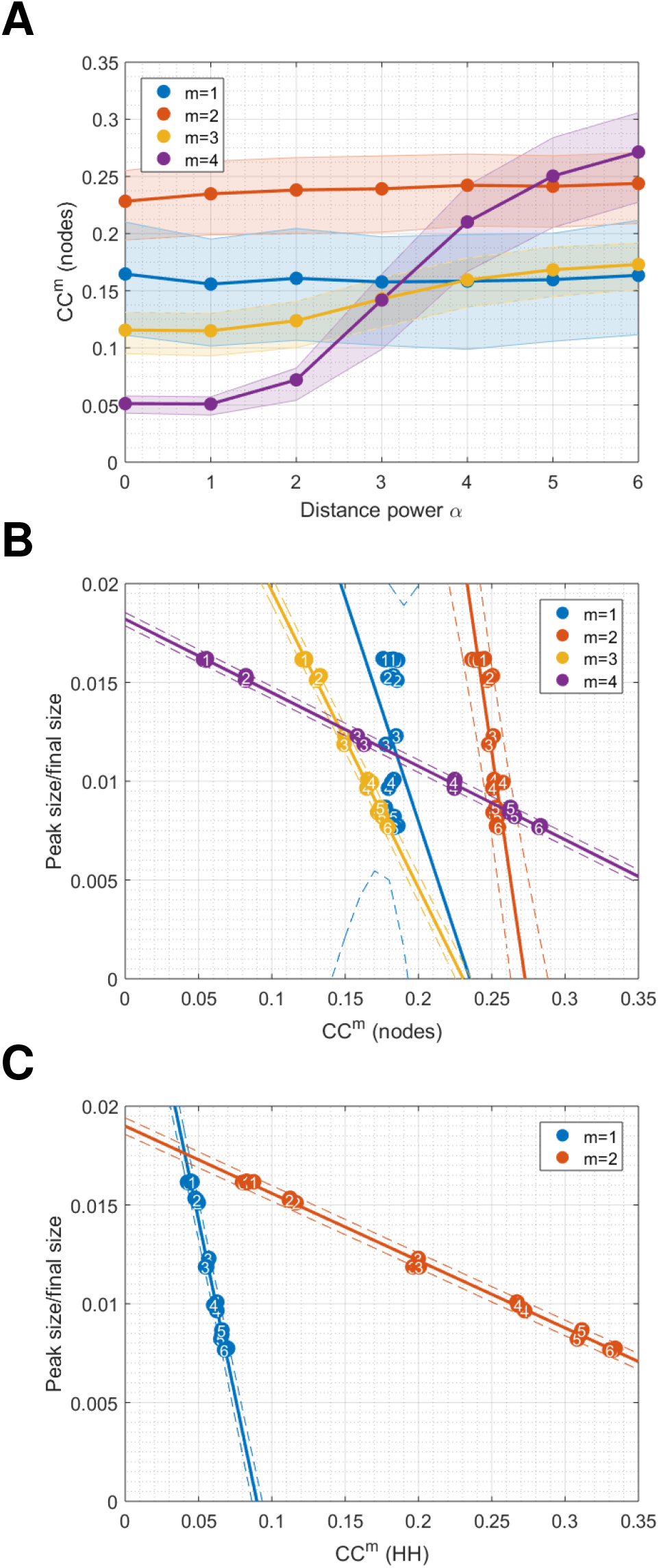
**(A)** 25−, 50− and 75-percentiles of order-*m* clustering *CC^m^* on networks constructed with different values of *α* and *h* = 5, *w* = 50, *p_w_* = 0.14, ⟨*k*⟩ = 10 and *R*_0_ ∈ [2, 2.2). Plot shows mean values over 3 different networks for each parameter set; **(B)** Using peak size as a crude metric for sub-exponential growth (given a fixed range for *R*_0_), we see linear trends emerging with higher orders of clustering. Plot shows one point per network, with 3 networks generated for each parameter set, and the mean peak size over 10 independently simulated epidemics, All points are numbered with the corresponding value of *α*; **(C)** Similarly for the household-only networks. Solid lines show linear fits to data, and dotted lines show 95% confidence intervals. Values of linear correlation coefficient and gradient of fits are given in SI Appendix, Table S2.

We conducted a number of sensitivity analyses for these network simulation results. Analytic approximations for degree distribution *P* (*K* = *k*) and expected first order clustering ⟨*CC*^1^⟩ in our networks are given in Protocol S1, and are independent of *α*. We confirmed these relationships in SI Appendix Figure S6 by computing these quantities on a set of networks that differ in *α*. SI Appendix Figure S7 shows the relationship between *α* and clustering order 1 to 4 on networks generated using a uniform population density. SI Appendix Figure S8 shows the relationship between order-*m* clustering *CC^m^* and peak size for different values of *R*_0_. SI Appendix Figure S 9 shows clustering order 1 to 4 on networks with different *h, w* and *p_w_*, and SI Appendix Figure S10 provides an illustration of the relationship between higher-order clustering and rewiring probability on a commonly used network model with spatial embedding: the Watts-Strogatz Small World Network [40].

Finally, we map our network model onto a deterministic metapopulation framework so as to relate our simulations of incidence to prior analytic approximations of travelling spatial waves (Protocol S1 for analytic construction). Figure 5 shows the results of simulating on a grid of evenly spaced households of size *h* = 4, where a single continuous variable describes prevalence in each household, and spatial coupling between households used in the force of infection is exactly the kernel used in the construction of our spatially embedded networks. We simulate with randomly spaced seeds (as above), and with a central seed (the center most 4 households), tracking global incidence and local time of peak incidence. The former case yields global incidence curves similar to those generated in our network model (which was seeded similarly). The latter case allows us to identify 4 distinct stages in the propagation of spatial waves that contribute to observed sub-exponential outbreak dynamics in more complex, network-based systems. SI Appendix Figure S 11 shows local peak timing in each case, and SI Appendix Figure S12 shows simulation results in 1 spatial dimension with *α* = 6 and *α* = 12, along-side statistical properties of prevalence, which further clarify these growth phases (c.f. figure captions for details and SI Appendix Protocol S1 for mathematical analysis).

**Figure 5:**
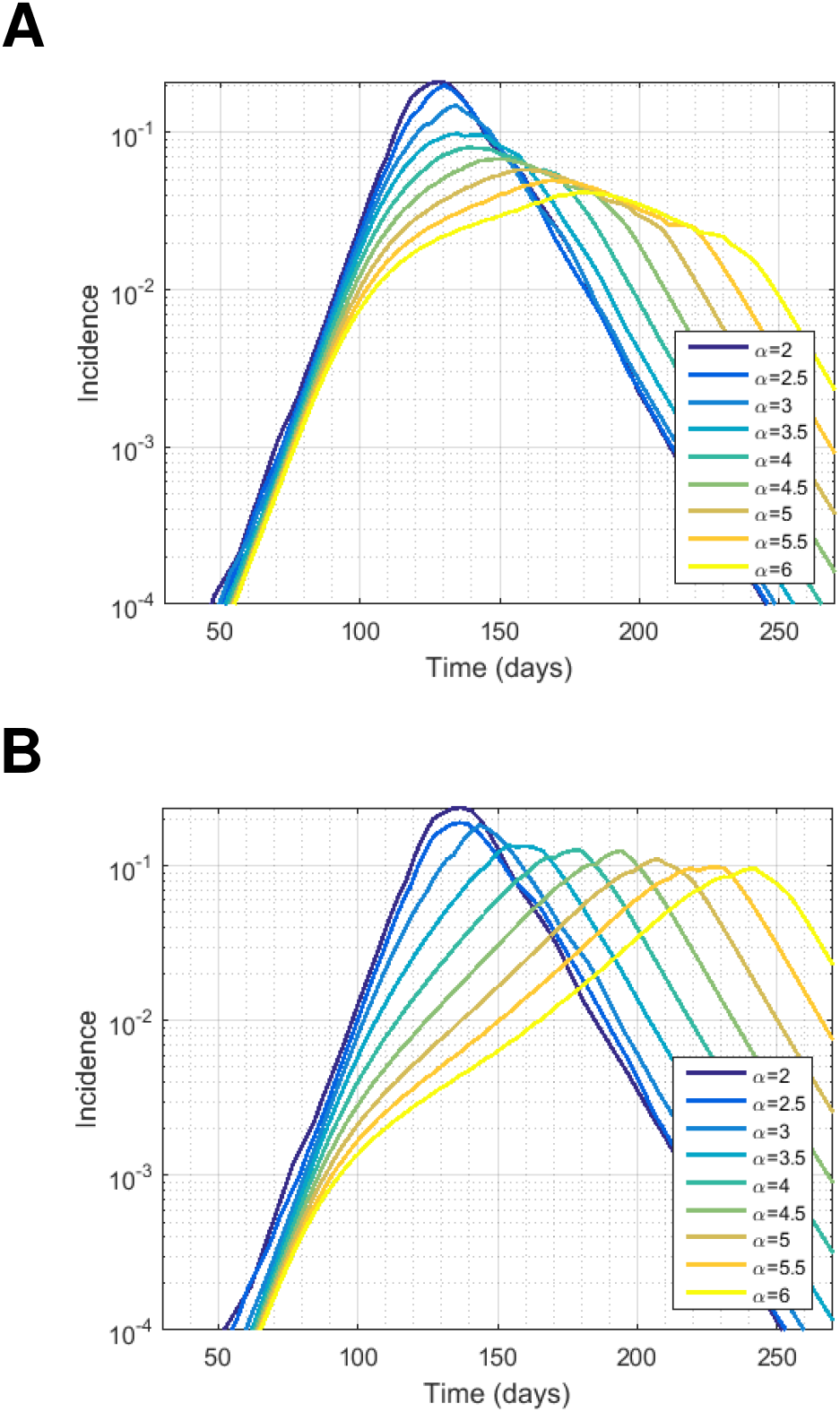
Mean-field approximation with *R*_0_ = 2.2, ⟨*k*⟩ = 10, *h* = 4, using a 100 × 100 grid of uniformly spaced households: **(A)** seeding in 10 randomly selected households (the same households are used in each simulation), and **(B)** seeding in the centre only. Incidence is given as a proportion of the total population for *α* ranging from 2 to 6. Supplementary Figure S10 shows time of peak incidence in the case *α* = 6 seeded as above.

## Discussion

We have shown that non-standard epidemic dynamics can arise from strongly spatially embedded social networks. Using a flexible algorithm of assigning individuals to households and then creating a social networks with widely varying topologies, we can explain the absence of exponential growth and increased attack rate for a given peak height in terms of higher order social structure, while maintaining a standard low-variance offspring distribution. We observe consistent patterns when we control for the basic reproductive number, as measured as directly as possible from a constant ratio of incidence between generations.

The algorithm we used [35] captures the key social contexts of home and workplace while using few parameters, which has allowed us to isolate specific relationships within the epidemic dynamics, across a broad range of network topologies. However, its simplicity is a potential limitation. Specifically, an individual only belongs to a single workplace (which may represent a school or social club). In reality, people will gather non-household contacts from a variety of sources. Also, our networks are not dynamic, which may limit the generalisability of the results to short generation time pathogens.

Accurate empirical data about higher order social contacts would allow us to address some of these issues. There are a number of different approaches to gathering social contact data, including contact diaries, mobile phone apps and tag-based location tracking [31]. Diary methods and current analytical approaches can provide accurate estimates of 1st order moments (degree distribution [32]) and valuable insights into second order moments (clustering [44]). However, these data and current analytical approaches are limited for the estimation of higher order moments. It seems likely that either high resolution mobile phone location data [7] or very high coverage tag-based studies will be needed to reveal these patterns [6]. In addition, further work is needed on the use of algorithms similar to that used here to explicitly fit fully enumerated social networks to egocentric sample data from a subset of the population (or low coverage non-egocentric data) [23].

Our results can be compared with other disease-dynamic models that produce non-standard incidence profiles. Different functional forms have been suggested for the force-of-infection term in compartmental models that give polynomial growth in the early stages of an epidemic [8, 18]. However, the key features of these model structures may be captured by a more straightforward underlying process [20]. Faster than exponential growth can be achieved with very high variance offspring distributions, which have been inferred by diary studies of social contacts [25]. There is also an extensive literature of much more abstract grid-based models of infectious disease that produce non-standard epidemic dynamic because of very local spatial processes (cellular automata [41]). We note that short periods of super-exponential growth were observed in our results for the simplified 2 dimensional metapopulation example (Fig 5B), arising from from accelerating spatial waves of incidence, not driven by the variance of the offspring distribution.

Prospective forecasting of infectious disease incidence during outbreaks [29] and seasonal epidemics [2] is an active area of public health research. Although non-mechanistic [13] and simple compartmental models [33, 39] have proven most reliable up to now, modern computing capacity enables studies to explore the possibility that incidence forecasts can be improved by the incorporation of realistic social network topology [36, 9]. For example, incidence of Ebola in west Africa in 2013-2016 and currently in central Africa exhibits strong spatial clustering and highly non-standard incidence dynamic, with short periods of exponential growth followed by low sustained peaks in incidence [10]. Future forecasting studies should explore the possibility that that sparse population density and short distances between contacts result in higher-order clustering in the social networks and the resulting non-standard incidence profiles.

## Methods

### The Model

We simulate 10 independent epidemics for each of 200 parameter sets (*h, v, p*_*w*_, *R*^∗^) drawn from a Latin hypercube, each seeded in 10 randomly selected individuals, and for each *α* = 0, 3, 6. The ranges of values used in the Latin hypercube are given in SI Appendix, Table S1, and complete parameter sets for all networks are given in SI Appendix, Table S1. Our simulations allowed us to track disease incidence and disease generation of each infection.

We simulate an epidemic on the network to reflect the natural history of Ebola, with a latent period of 9.7 days and a serial interval of 15.3 days. The generation time was calibrated by varying the relative infectiousness of a short period before the onset of symptoms. Global transmissibility *β* is tuned to the value of *R*^∗^ drawn from the Latin hyper-cube. For each timestep, the probability of infection is calculated for each edge in the network. The algorithm progresses in real time with small timesteps so it can be compared with results from compartmental models. Details of the network simulation algorithm are given in [35].

### Assigning *R*_0_ to each simulation

For each simulation output, we calculate the mean reproductive ratio for each generation. For generations 1 to 9 and for each possible consecutive string of 3, 4 or 5 values, we perform a linear regression fit. We define *R*_0_ as mean reproductive ratio over the set of values for which the gradient of this fit is closest to 0 (and all values the remain larger than 1). This allows us to assign a value *R*_0_ to every simulation output.

### Higher order clustering

We compute our higher-order clustering coefficients on a subset of 1000 nodes in each network, chosen at random. The algorithm involves storing the network structure as lists of neighbours for each node, and performing an effective contact-tracing procedure. Though it is possible to compute these metrics for all nodes via successive multiplication of adjacency matrices, this procedure becomes computationally expensive in higher orders at networks become large.

## Data availability

Results can be reproduced in the Ebola scenario in the id_spatial_sim repository [5], using scripts ebola_build.sh and ebola_run.sh.

## Acknowledgements

We thank Derek Cummings for useful discussions. For funding, we thank UK Medical Research Council (MRC) and the UK Department for International Development (DFID) under the MRC/DFID Concordat agreement, also part of the EDCTP2 programme supported by the European Union (UK, Centre MR/R015600/1) (DH & SR); Wellcome Trust Investigator Award (UK, 200861/Z/16/Z) (SR); Wellcome Trust Collaborator Award (UK, 200187/Z/15/Z) (SR); National Institute for General Medical Sciences (US, MIDAS U01 GM110721-01) (DH,SR); National Institute for Health Research (UK, for Health Protection Research Unit funding) (SR); and the Center for Disease Control (DH).

