## Supplementary Information for "Social networks with strong spatial embedding generate non-standard epidemic dynamics driven by higher-order clustering"

### Protocol S1: analytic approximations

#### Derivation of degree distribution and global clustering coefficient

$$\begin{aligned}
\langle k\rangle &= h-1+(w-1-\langle housemates\ at\ workplace\rangle)p_w \\
&\approx h-1+p_w(w-1) \\
P(K=k) &= \sum_{1\le i\le k+1} P(H=i) \sum_{j-1\ge k-i+1} P(W=j) \\
&\times \binom{j-1}{k-i+1} p_w^{k-i+1}(1-p_w)^{j-1-k+(i-1)} \\
&= \sum_{1\le i\le k+1} \frac{(h-1)^{i-1}e^{1-h}}{(i-1)!} \sum_{j\ge k-i+2} \frac{(w-1)^{j-1}e^{1-w}}{(j-1)!} \\
&\times \binom{j-1}{k-i+1} p_w^{k-i+1}(1-p_w)^{j-k+i-2} \\
\langle CC^1\rangle &\approx \frac{(h-1)(h-2)+p_w[p_w(w-1)-1][p_w(w-1)-2]}{\langle k\rangle(\langle k\rangle-1)}
\end{aligned}$$

where denominator $= \frac{1}{2}E(potential\ contacts\ between\ neighbours)$ and numerator $\approx \frac{1}{2}E(actual\ contacts$

#### Metapopulation approximation

We now consider a metapopulation approximation to our network model, within the framework of renewal equations. This will allow us to use existing theory in order to interpret our results.

Let $i, j, k$ index the households in a given population, $h$ the mean household size and $\langle k\rangle$ the mean contact rate of an individual (this notation is consistent with mean degree the corresponding network model). We let $K_{ij}$ denote he connection density between households such that $K_{ii} = 0\ \forall i$ and $\sum_j K_{ij} = 1\ \forall j$, and we define the workplace contact rate $v = \langle k\rangle - h + 1 = p_W(w-1)$.

The expected force of infection on an individual in household $i$ (denoted $[i]$) is then given by:

$$\langle \lambda_{[i]}\rangle = \frac{\beta}{\langle k\rangle}\left[I_i + \frac{v}{h}\sum_j K_{ij}I_j\right] \tag{1}$$

where the denominator $h$ comes from the fact that $\sum_j K_{ij}\langle N_j\rangle = h$, where $\langle N_j\rangle = h$ denotes the expected population of household $j$.

The parameter $\hat{\rho} := h/v$ controls the diagonal weighting of the global coupling kernel $E + \hat{\rho}K$, where $E$ denotes the $n \times n$ identity matrix (avoiding confusion with $I$, which is reserved for infectious populations). Furthermore, our spatial coupling parameter $\alpha$ appears explicitly in $K$:

$$K_{ij} \propto \frac{1}{1+(d_{ij}/a)^{\alpha}} \tag{2}$$

Note that, though $K$ is typically called a "travel kernel", we are describing a static network structure with no (explicit) travel. Also, the household/workplace framework does not lend itself to a description in continuous space, since households correspond to single points in which finitely many agents are densely connected. We can, however, proceed with one continuous variable per household and, for any given population density, we can define a discrete space in which each location contains (at most) one household.

The presence of local decay in infectives *whilst the global epidemic is still growing* also renders the system described in Equation (1) somewhat intractable analytically. However, if we view the force of infection $\langle\lambda_{[i]}\rangle$ as follows:

$$\langle\lambda_{[i]}\rangle = \langle\lambda_{[i]}^{in}\rangle + \langle\lambda_{[i]}^{out}\rangle \tag{3}$$

$$= \frac{\beta}{\langle k\rangle} I_i + \frac{\beta}{\hat{\rho}\langle k\rangle}\sum_j K_{ij} I_j \tag{4}$$

then the limiting cases $v \to 0$ and $h \to 1$ provide additional insight into the different growth phases observed. The first growth phase is simply the mass-action SIR model, and the final, accelerating wave-like phase is related to the continuous-space model defined by:

$$\lambda(t;\mathbf{x}) = \tilde{\beta}\frac{\int I(t;\mathbf{y})K(\mathbf{x}-\mathbf{y})d\mathbf{y}}{\int N(\mathbf{z})K(\mathbf{x}-\mathbf{z})d\mathbf{z}} \tag{5}$$

Moreover, both of these limiting cases are covered substantially in the literature. In particular, we draw the reader's attention to the continuous space case, in which radial wave solutions are known to exist for $\alpha > 3$, with constant speed for $\alpha > 4$ and accelerating for $\alpha \in (3, 4]$ [1, 4, 2, 3].

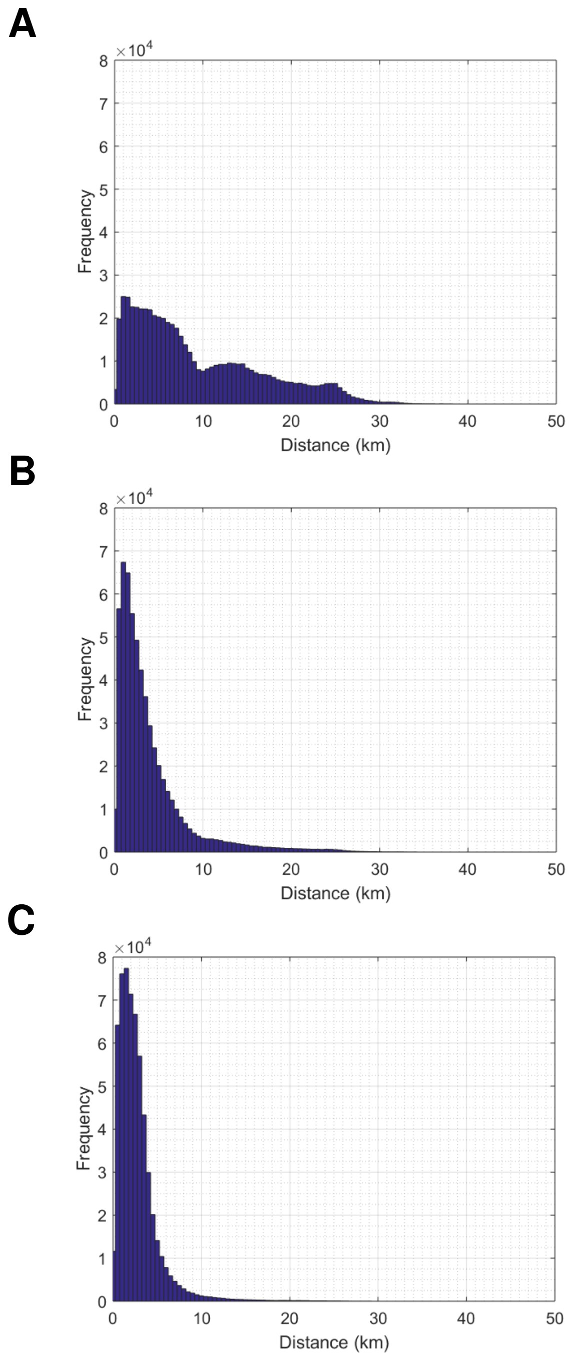


Figure S1: Examples of commuting distributions for **(A)** $\alpha = 0$, **(B)** $\alpha = 3$ and **(C)** $\alpha = 6$. All networks use population of Monrovia, with $h = 4,\ w = 50$ and $p_w = 0.14$, so $\langle k \rangle = 10$.

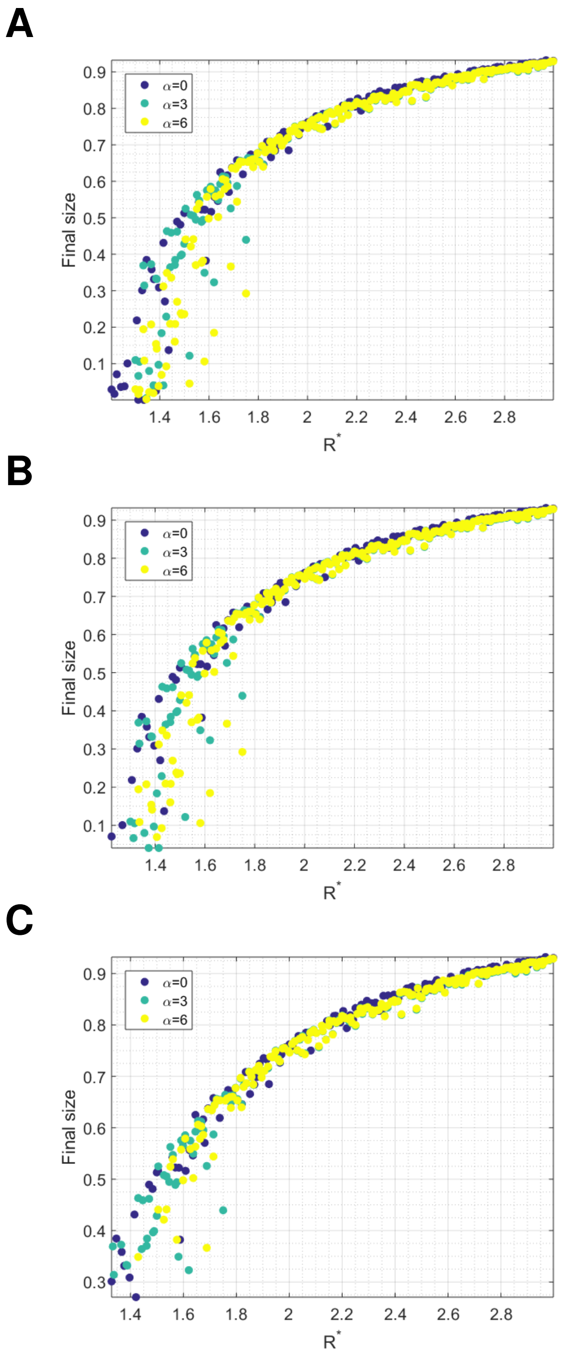


Figure S2: The relationship between $R^*$ and final size for **(A)** all simulations **(B)** all with peak over $100$ and **(C)** peak over $500$ (all simulations have seed size $10$).

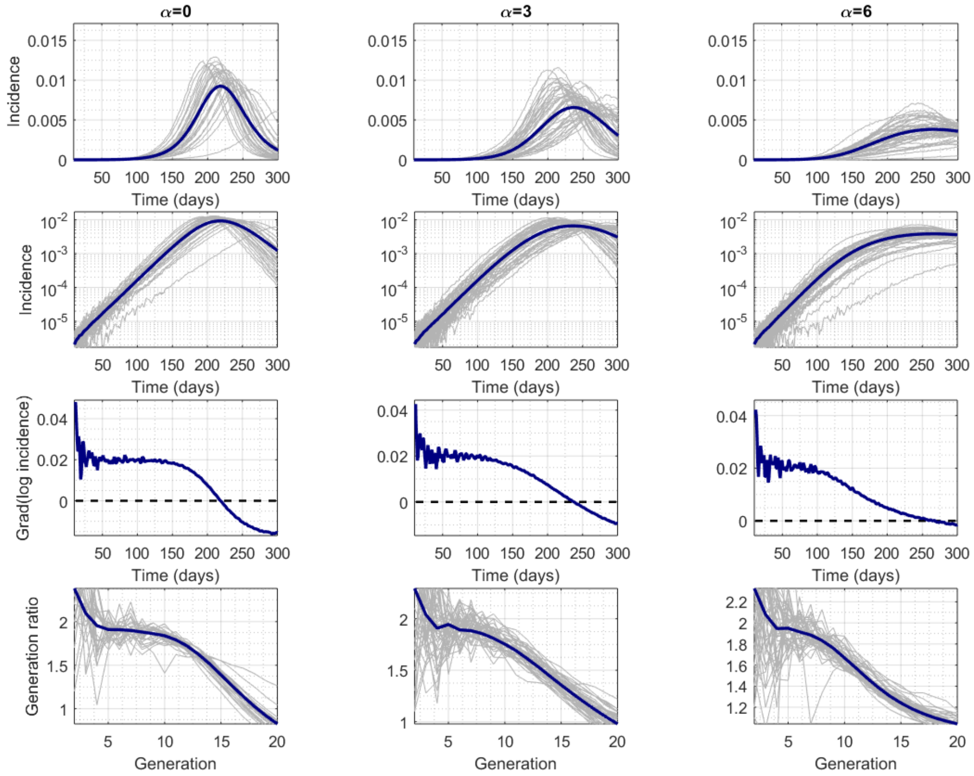


Figure S3: Simulation results for $R_0 \in [1.8, 2),\ \alpha = 0, 3, 6$.

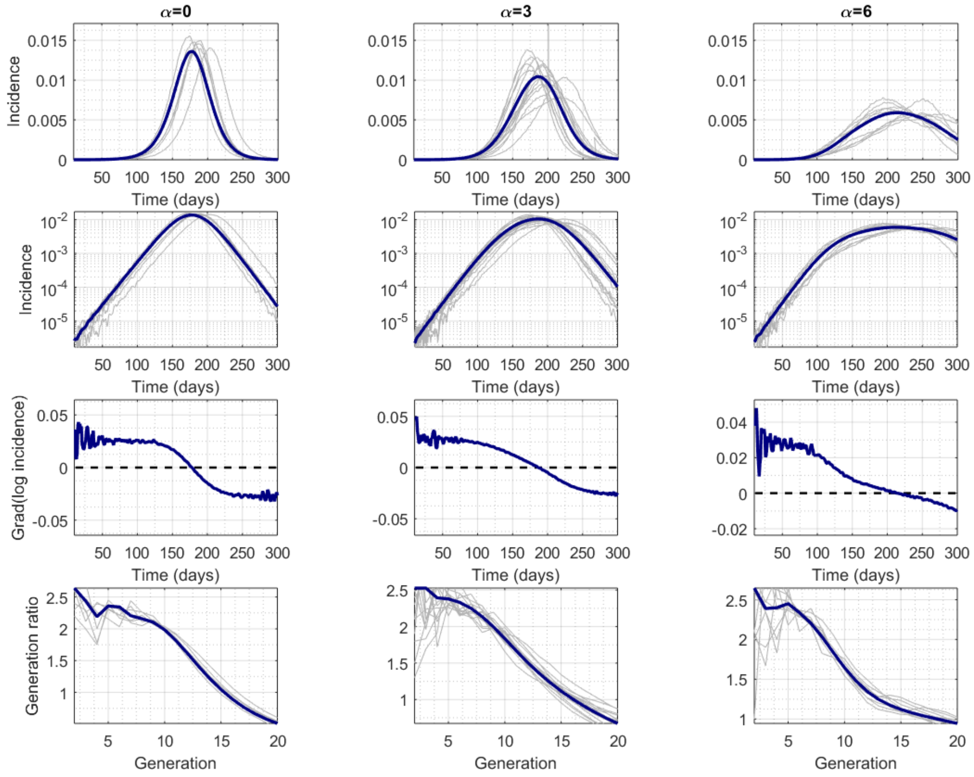


Figure S4: Simulation results for $R_0 \in [2.2, 2.4),\ \alpha = 0, 3, 6$.

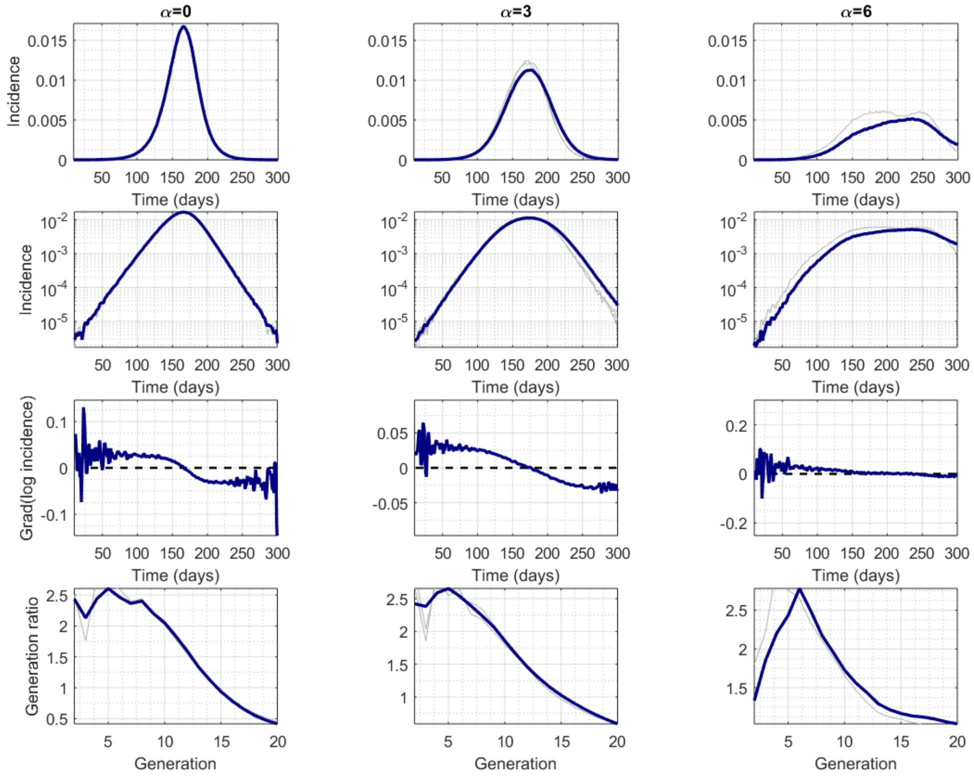


Figure S5: Simulation results for $R_0 \in [2.4, 2.6),\ \alpha = 0, 3, 6.$

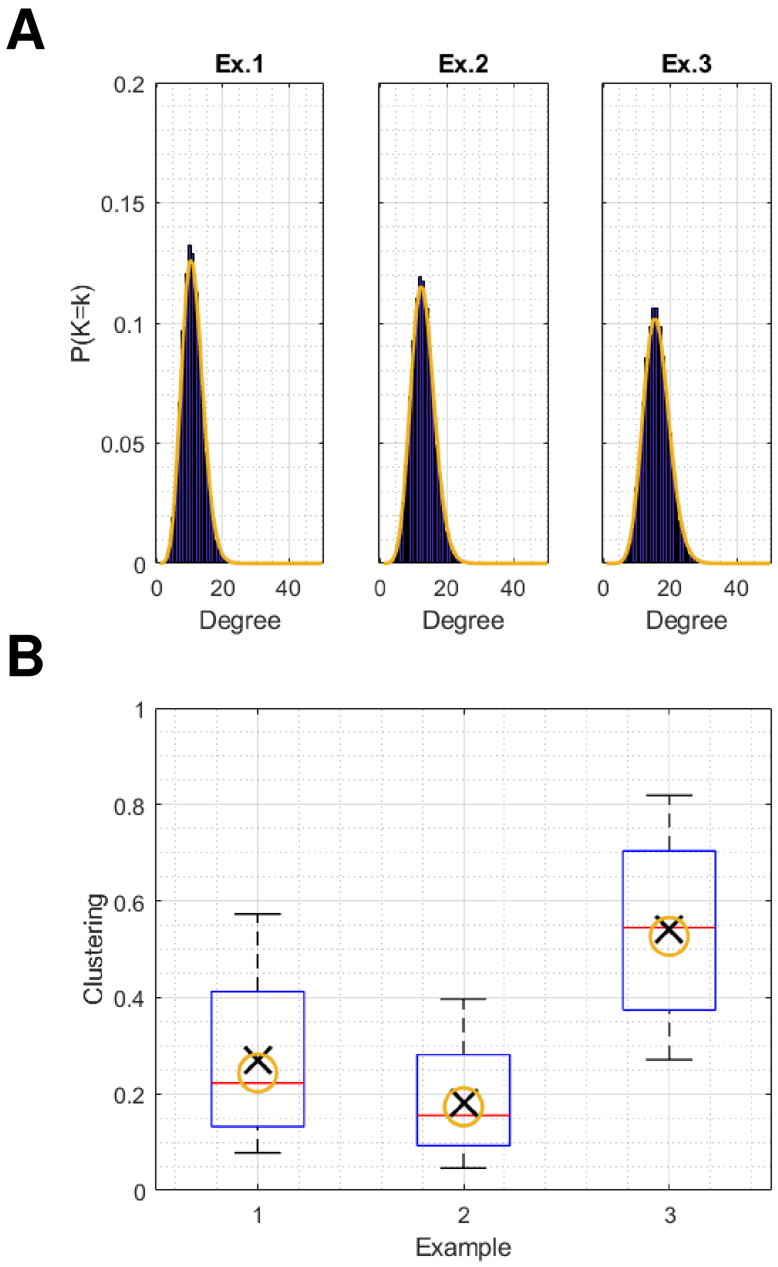


Figure S6: **(A)** Degree distribution, and **(B)** clustering coefficient for 3 example networks, with analytic approximations for $P(K=k)$ and $\langle CC^1 \rangle$ shown in yellow. Example 1: $h=6,\ w=50,\ p_w=0.1,\ v=5,\ \langle k \rangle = 10$; example 2: $h=6,\ w=100,\ p_w=0.07,\ v=,\ \langle k \rangle =$, example 1: $h=11,\ w=80,\ p_w=0.05,\ v=4,\ \langle k \rangle = 14$.

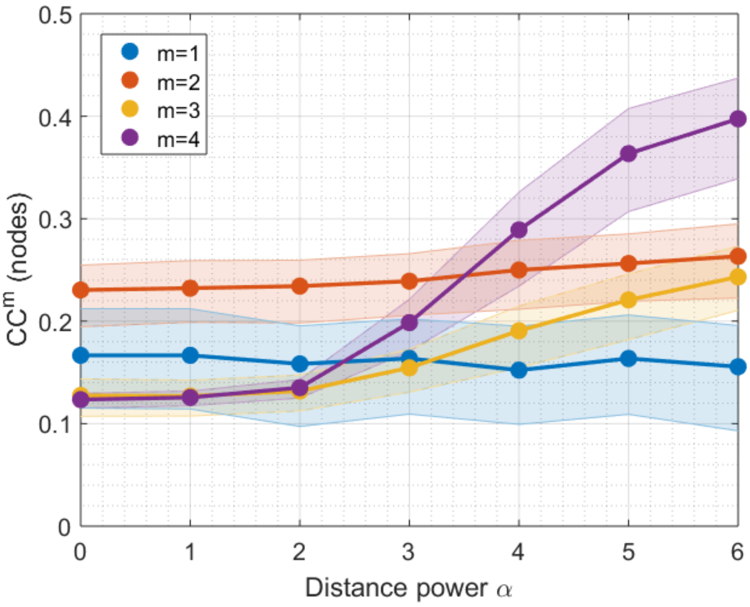


Figure S7: Relationship between spatial correlation parameter $\alpha$ and clustering order $1$ to $4$ on a uniform population, defined as a $50 \times 50,\ 1\text{km}^2$ grid with a population of $20$ per pixel, and $h = 5,\ w = 50,\ p_w = 0.14,\ v = 7,\ \langle k \rangle = 11$ as in main Figure 4.

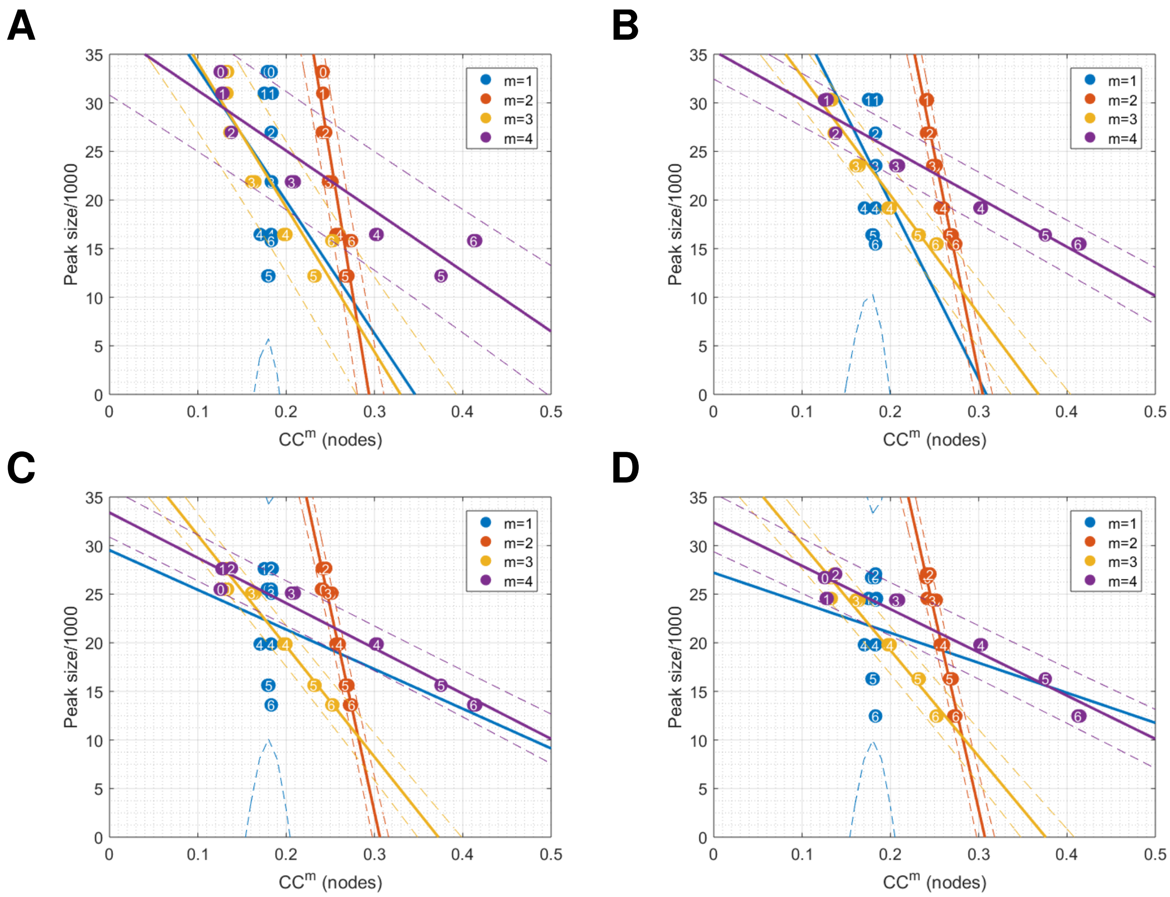


Figure S8: Relationship between order-$m$ clustering an epidemic peak height on the uniform population defined in Figure S7: **(A)** $R_0 \in [1.8, 2)$, **(A)** $R_0 \in [2, 2.2)$, **(A)** $R_0 \in [2.2, 2.4)$, **(A)** $R_0 \in [2.4, 2.6)$.

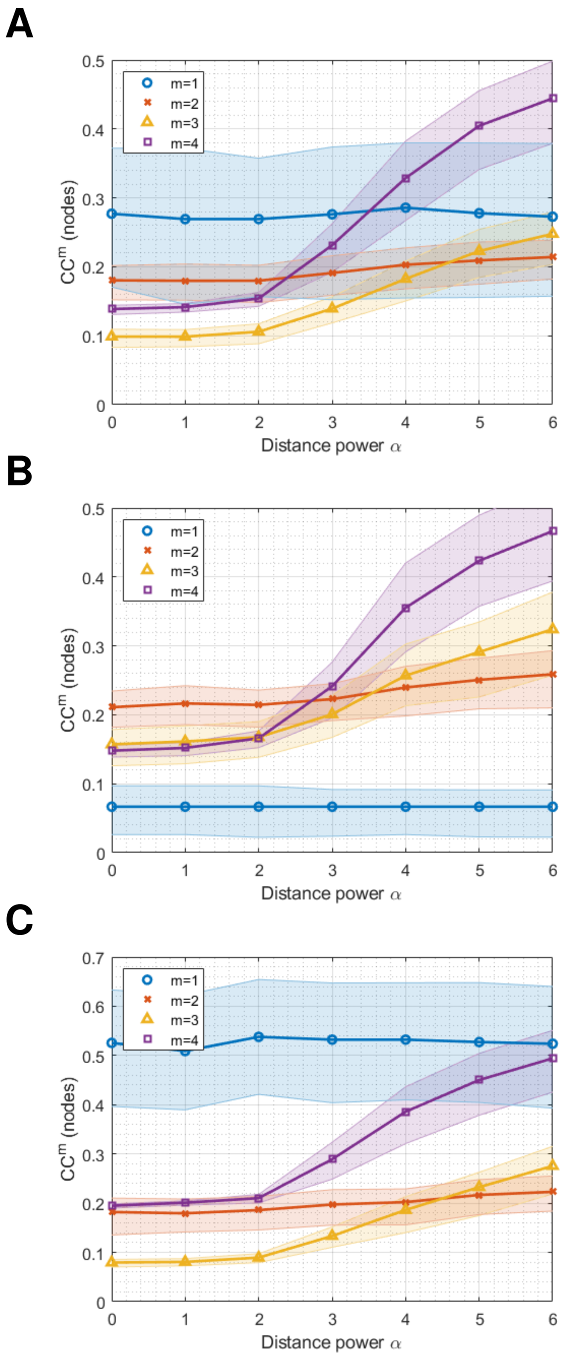


Figure S9: Sensitivity analysis: clustering orders 1 TO 4 for $\alpha = 0, 1, ..., 7$ for a sample of 1000 nodes on networks with **(A)** $h = 6, w = 100, p = 0.05$; **(B)** $h = 3, w = 200, p = 0.04$; **(C)** $h = 12, w = 100, p = 0.04$.

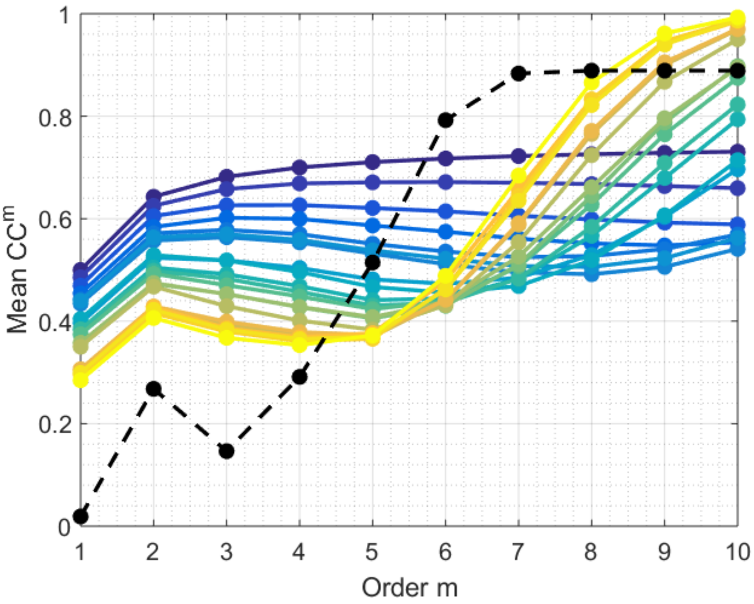


Figure S10: Illustration of higher order clustering in Watts-Strogatz Small World networks $G(n, m, p)$. We fix number of nodes $n = 10^3$ and mean degree $k = 4$. Rewiring probability ranges from $0$ (blue) to $0.2$ (yellow). The black dotted line shows values for random graphs ($p = 1$).

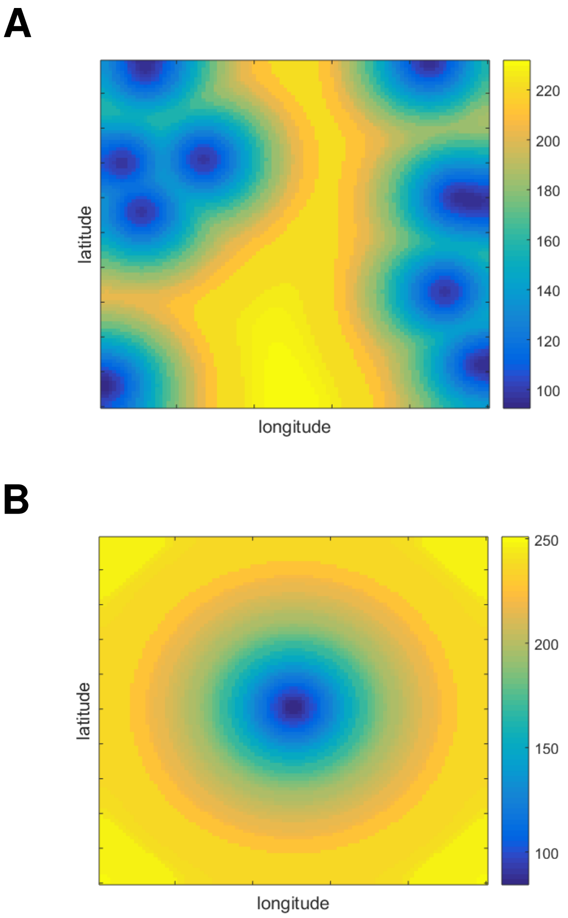


Figure S11: Peak times for $\alpha = 6$: **(A)** seeding in 10 random locations and **(B)** seeding in the centre.

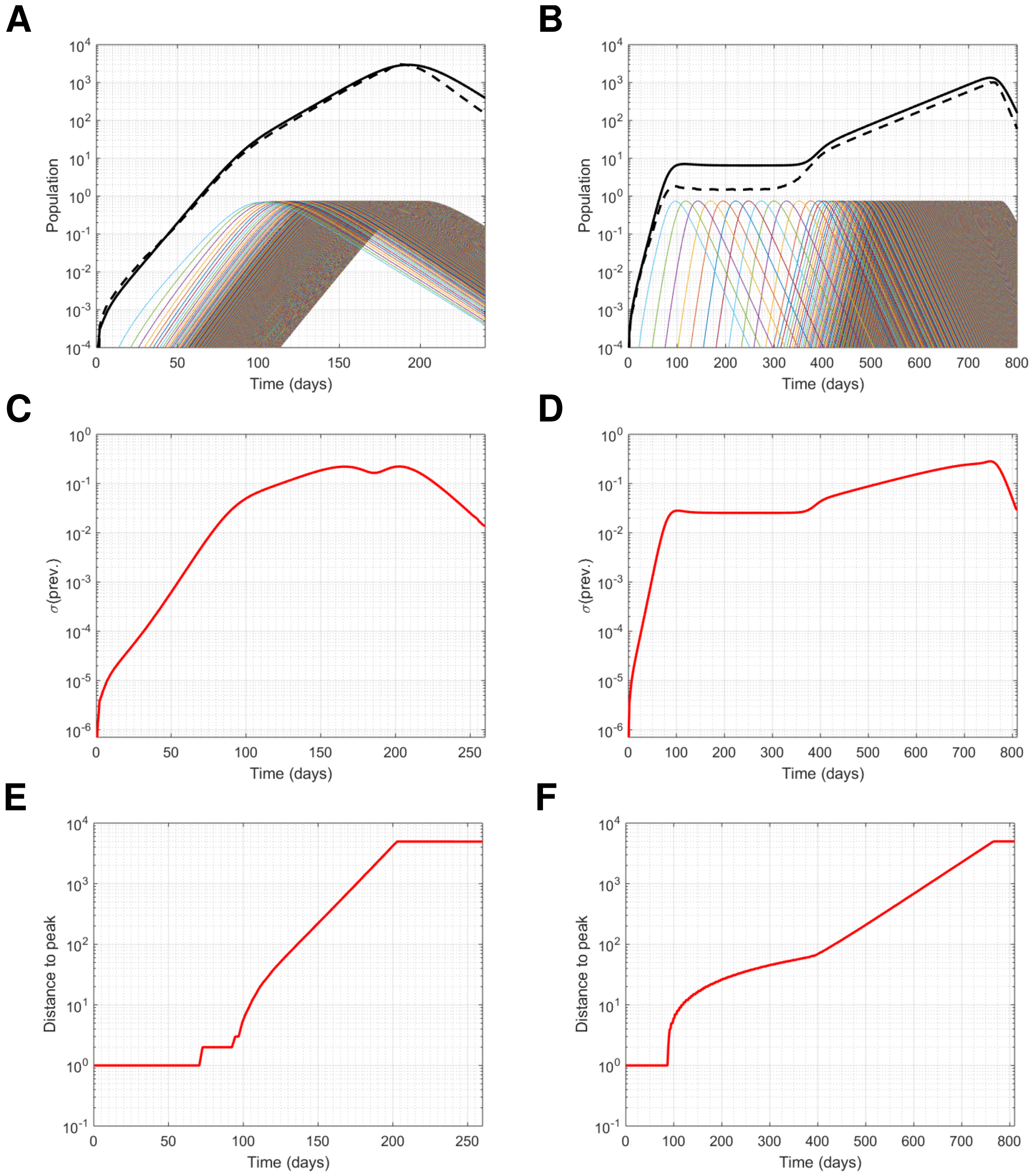


Figure S12: Mean-field approximation with $\langle k \rangle = 10,\ h = 4$, using a $1 \times 5000$ grid (1D model) of uniformly spaced households and seeding at one end: **(A)** global prevalence (solid black line), global incidence (dashed line) and local prevalence in every $5$th location (colored lines), **(C)**, standard deviation of prevalence, **(E)** distance to peak prevalence for $\alpha = 2$, **(B)**, **(D)**, **(F)** the same with $\alpha = 10$. Though unrealistic, using $\alpha = 10$ in $1$ spatial dimension shows $4$ clear, distinct growth phases: $1$. within-seed-household exponential growth, $2$. constant-speed travelling wave, $3$. distribution of infectives gets broader as enough within-household infections overlap in time, and $4$. accelerating wave-like solution (though s.d. of spatial distribution also grows exponentially). Comparing with $\alpha = 3, \ldots, 6$ and different seeding patterns in 2D (c.f. main Figure 5), we see how these $4$ phases overlap to give sub-exponential growth.

#### Supplementary Tables

| Parameter | Min | Max |
|---|---|---|
| $h$ | 2 | 8 |
| $v = wp_w$ | 5 | 20 |
| $p_w$ | 0.1 | 0.8 |
| $R^*$ | 1.3 | 3 |

Table S1: Ranges of each parameter value in the Latin hypercube.

| Order | Corr. | Grad. |
|---|---|---|
| 1 | 0.2599 | 0.1131 |
| 2 | -0.7538 | -0.4082 |
| 3 | -0.996 | -0.134 |
| 4 | -0.9972 | -0.0163 |
| 2 (HH) | -0.9208 | -0.2828 |
| 4 (HH) | -0.9908 | -0.0169 |

Table S2: Linear correlation coefficients and gradients of linear regression model for main Figures 4B (nodes) and 4C (households).
